# Development and validation of allele specific *RLK* gene markers for banana bunchy top disease (BBTD) resistance in banana

**DOI:** 10.1101/2023.12.20.572701

**Authors:** Reina Esther S. Caro, Anand Noel C. Manohar, Roanne R. Gardoce, Fe M. Dela Cueva, Lavernee S. Gueco, Ma. Carmina C. Manuel, Darlon V. Lantican

## Abstract

As global demand for bananas rises, the threat of banana bunchy top disease (BBTD), caused by *banana bunchy top virus* (BBTV), poses a significant constraint to sustainable banana production. To circumvent this, it is critical to fast-track the development of BBTV-resistant banana cultivars. In a dataset of differentially expressed genes between BBTV-resistant and susceptible bananas, six *RLK* genes involved in plant defense against pathogens were exclusively upregulated in the BBTV-resistant wild *Musa balbisiana* compared to the BBTV-susceptible *Musa acuminata* cv. ’Lakatan’. Here, two genic variants of *RLKs*, single nucleotide polymorphisms (SNPs) and simple sequence repeats (SSR), were identified in 10 banana accessions representing major genomic groupings. Nine gene-specific markers were developed using three marker systems: SSR, allele-specific PCR, and tetra primer amplification refractory mutation system PCR markers. Six of these were polymorphic and successfully discriminated 117 *Musa* genotypes at the NPGRL-IPB *ex situ* germplasm, with an average polymorphism information content of 0.36 for SNP markers and 0.49 for SSR marker, both of which are moderately informative. These markers facilitated the construction of a dendrogram using the Unweighted Neighbor-Joining (UNJ) method with 1000 bootstrap iterations, identifying four major clades: *M. acuminata* spp. and other *Musa* spp. (I), wild *M. balbisiana* (BBw) and other accessions with high-proportion of B-genome (II & III), hybrid bananas (triploids and tetraploids) (IV), which revealed the genetic diversity and relationships among *Musa* accessions. Ultimately, this genomic resource will aid in positioning the Philippines as a key player in banana disease resistance breeding towards a food-secure banana industry.

## INTRODUCTION

Bananas are considered to be one of the most highly valuable global crops. In 2020, it solidified its position as the top traded fruit worldwide, underlining its immense value in the global agricultural industry. With an estimated total production of 120 million tons (FAO 2022), banana stands tall as one of the world’s most economically influential commodities. Among the major key players in banana production, India takes the lead with 26.3% contribution to the total global production, following closely behind are countries such as China, Indonesia, Brazil, Ecuador, and the Philippines (FAOSTAT 2020), which further substantiates the significance of this crop across different economies. In the Philippines, bananas are the leading fruit crop and country’s top agricultural export commodity in 2019, with an export volume of 4.4 million metric tons valued at Php 101.18 billion (PSA 2019). With this, the Philippines emerged as one of the significant players in the banana export industry, consistently ranking as the second-largest banana exporter in the recent years. However, due to the ongoing production challenges, it has recently dropped to the third place next to Ecuador and Guatemala in the global banana exports in 2022 (FAO 2022).

Bananas are classified within the Musaceae family under the genus *Musa* (Simmonds 1995; Pachuau et al. 2014). They have originated in Southeast Asia and has been distributed and cultivated in more than 130 tropical and subtropical countries, and western Pacific countries (Govaerts and Häkkinen 2006; Resmi and Nair 2007; Davey et al. 2013; Janssens et al. 2016). Edible banana cultivars were derived from two (2) wild seeded diploid species of *Musa acuminata* Colla (AA) and *Musa balbisiana* Colla (BB) (De Langhe et al. 2009; Perrier et al. 2011; Martin et al. 2020) and diversified into different genome combination with various ploidy level such as diploids (AA; BB; or AB; 2n = 2X = 22); triploids (AAA; AAB; or ABB; 2n = 3X = 33); and tetraploids (AAAA; AAAB; AABB; or ABBB; 2n = 4X = 44) (Simmonds and and Shepherd 1955). Commercially available banana cultivars are often triploid hybrids and are propagated vegetatively due to high levels of sterility (Raboin et al. 2005; Perrier 2009; Perrier 2011; Ortiz and Swennen 2014).

Beyond their nutritional benefits and delightful taste, bananas have fostered job opportunities, contributed to economic growth, and supported sustainable development in various countries, highlighting the banana’s potential as a catalyst for progress and prosperity within the agricultural sector. However, despite their economic importance, bananas are susceptible to a range of catastrophic diseases, one of which is the banana bunchy top disease (BBTD) caused by *banana bunchy top virus* (BBTV), vectored by banana aphids (*Pentalonia nigronervosa*), infecting various members of Musaceae like bananas, plantains, abaca, and ornamental bananas in a persistent circulative and non-replicative manner (Harding et al. 1991; Hu et al. 1996; Hong-Ji 2000; Fauquet et al. 2005; Robson et al. 2006; Qazi 2015; Dale et al. 2017). The disease is also easily spread by the use and exchange of infected planting materials or vegetative propagules such as corms and suckers among banana growers (Thomas et al. 2003). BBTV infected plants showed systematic symptoms such as stunted growth, rosetted appearance, and chlorotic leaves affecting the production of harvestable fruit bunches (Molina et al. 2009).

BBTD has been continuously spreading in different banana producing countries, mostly in the Asia Pacific region, which contributed up to 100% yield loss in the banana production in severely infected and highly susceptible plants and caused a drastic reduction in the cropping area (Kumar et al. 2015). In the Philippines, BBTD incidence has been slowly increasing for the past decades which greatly affected the livelihood of smallholder farmers as well as large-scale growers (Molina et al. 2009). Controlling the spread of disease in bananas includes elimination of infected plants, use of virus-free planting materials, management of insect vector populations, and early detection of infected plants (Hong-Ji 2000; Bajet and Magnaye 2002; Dela Cueva et al. 2010; Arumugam et al. 2017). However, the use of BBTV-resistant banana cultivars remain to be the most effective and sustainable way of managing disease and minimizing yield loss in banana production (Jekayinoluwa et al. 2020; Tripathi et al. 2020).

Early reports have shown that there are currently no natural sources of BBTV resistance, but some banana cultivars with a component of the B-genome derived from wild *M. balbisiana* (BBw) have demonstrated a level of disease resistance compared to those with purely A genome composition (Hapsari and Masrum 2012; Ngatat et al. 2017; Latifah et al. 2021). Resistance evaluations in the Philippines, however, have identified wild *M. balbisiana* accessions to have complete resistance or immunity to the virus (Dela Cueva et al. 2023). Utilizing these germplasm materials, Lantican et al. (2023) identified differentially expressed genes in two distinct banana genotypes: BBTV-resistant wild *M. balbisiana* (BBw) and susceptible *M. acuminata* cv. ‘Lakatan’ (AAA), making a significant stride in understanding the molecular basis of banana resistance against BBTV. Interestingly, the study discovered a set of receptor-like kinases (*RLKs*), which are the first line of plant defenses against pathogens, exclusively upregulated in the wild *M. balbisiana* upon introduction of BBTV.

Plant *RLKs* function as pattern recognition receptors (PRRs) that detect pathogen-derived molecular patterns activating plant’s defense mechanisms during plant-pathogen interaction (Boller and Felix 2009). While *RLKs* have been studied in various plant species such as Arabidopsis (Shiu et al. 2004; Jose et al. 2020), soybean (Liu et al. 2009), wheat (Yan et al. 2023), and rice (Song et al. 1995; Shiu et al. 2004), their specific involvement in banana’s resistance to BBTV remains unexplored. By further investigating the differentially expressed banana *RLK* genes identified by Lantican et al. (2023) using complementary approaches (i.e., genic variation identification, wet laboratory investigations, and genetic association studies), researchers can strengthen our understanding on their potential roles in defense mechanisms against BBTV infection and uncover other genetic factors influencing BBTD resistance of bananas.

In this work, we generally aim to develop polymorphic DNA markers tagging putative *RLK* genes potentially associated with BBTV resistance in banana. This entailed identifying the genic variants of *RLKs* across a carefully selected panel of banana accessions from which polymorphic DNA markers were developed and applied across selected banana accessions. In addition, these markers were used to assess genetic diversity and relationships of banana germplasm collections, validate DNA marker’s efficiency and applicability in *ex-situ* germplasm management, and to establish association of the markers to BBTV infection in natural environment towards marker-assisted plant breeding applications. Ultimately, the development of gene-specific markers in these *RLK* genes will help in fast-tracking BBTV resistance breeding in banana.

## MATERIALS AND METHODS

### Biological sample preparation

Ten banana accessions representing different major banana genomic groups were carefully selected from the 183 banana accessions in the *ex situ* germplasm collection of the National Plant Genetic Resources Laboratory at the Institute of Plant Breeding (NPGRL-IPB)-UPLB utilized in the study of Gardoce et al. (2023). These accessions were utilized as plant materials to perform partial gene sequencing of putative *RLKs* as the basis for identification of potential polymorphic regions as target for DNA marker development (Table 1).

**Table 1.** List of ten (10) banana accessions selected for partial sequencing of six *RLK* genes.

| Entry # | Accession name | Genome composition |
| --- | --- | --- |
| 1 | Zebrina | AA |
| 2 | Pisang Keling | AAB |
| 3 | Lakatan Cavite 04-077 | AAA |
| 4 | Fhia 1 01 026 | AAAB |
| 5 | Latundan Davao | AAB |
| 6 | Kalapagnon 09-098 | ABB |
| 7 | Daydayawen | ABBB |
| 8 | Saba 10-049 | BBB |
| 9 | Moko | BBw |
| 10 | Cantong | Other (presumed as B × T) |

### DNA extraction and quantification

Genomic DNA was extracted from the second young leaf tissue of the selected banana accessions using DNA extraction protocol adapted from Doyle and Doyle (1990) with modifications. Extracted DNA samples were quantified using lambda (λ) DNA molecular weight standards (Sigma-Aldrich Inc., St. Louis, Missouri, USA) of known concentrations in a 1% UltraPure™ agarose gel (Invitrogen Corp., Carlsbad, California, USA) by densitometric means. DNA working stock (10 ng/uL) solutions were prepared and stored at -20°C.

### Partial gene sequencing of banana *RLK* genes

Six (6) upregulated receptor-like kinase (*RLK*) genes exclusive to wild *M. balbisiana* (BBTV-resistant) were obtained from the study of Lantican et al. (2023), as summarized in Table 2, and their corresponding nucleotide sequences were retrieved from the Banana Genome Hub (https://banana-genome-hub.southgreen.fr/) platform. Sequencing primers per gene were designed using SnapGene software (from Insightful Science; accessible at snapgene.com) following the general guidelines for PCR primer design: primer length of 17-24 bases, 40-50% GC content, minimum T_m_ mismatch of 2-3°C, G/C clamp at 3’ end, and avoidance of repeats and runs, as well as secondary structures (hairpin maximum ΔG: -0.3 kcal/mol; self-dimer maximum ΔG: -0.6 kcal/mol; and cross dimer maximum ΔG: -0.9 kcal/mol). Primer specificity and thermodynamic properties were assessed using Primer-BLAST (https://www.ncbi.nlm.nih.gov/tools/primer-blast) software and IDT OligoAnalyzer™ Tool (Integrated DNA Technologies, Inc., Coralville, Iowa; http://sg.idtdna.com/calc/analyzer). The amplicon size for each gene was restricted to less than 1.5 kb due to limitations in the sequencing platform’s chemistry. Prior to synthesis, designed primers were tested for *in silico* PCR across different established banana genomes in Banana Genome Hub (https://banana-genome-hub.southgreen.fr/) to discover gene regions that may differ among distinct *Musa* genomes. The designed sequencing primers (see Supplementary Table S1) targeting partial *RLK* gene regions were sent for outsourced oligonucleotide synthesis at SBS Genetech Co., Ltd. (Beijing, China). Optimized PCR reaction had a total volume of 10 uL consisting of 1X PCR buffer (10 mM Tris pH 9.1 at 20°C, 50 mM KCl, 0.01% Triton™ X-100, Vivantis Technologies, Malaysia), 2 mM MgC_l2_, 0.2 mM dNTPs (Promega Corporation, Madison, Wisconsin, USA), 0.1 uM each of forward and reverse primer (SBS Genetech Co., Ltd., Beijing, China), 0.5 U recombinant *Taq* DNA polymerase (Vivantis Technologies, Malaysia), and 10 ng DNA. The following PCR thermocycler settings were used: 94°C for 3 min, 35 cycles of 94°C for 30 sec (denaturation), 45-65°C for 45 sec (annealing), 72°C for 1 min (extension), and 72°C for 5 min for final extension. After primer optimization, a total of 30 ul PCR product per target gene were sent for outsourced double-pass capillary sequencing at 1st BASE (Apical Scientific, Malaysia) and raw sequences were subjected to standard sequence quality control procedures using Geneious software (Kearse et al. 2012). Sequences with percent high quality (%HQ) > 80 were selected and consensus sequences were generated for each *RLK* gene for subsequent analyses.

**Table 2.** List of upregulated receptor-like kinase (*RLK*) genes in BBTV-resistant wild *Musa balbisiana* accession (Lantican et al., 2023) and their functions.

| Entry # | Gene name | Function | References |
| --- | --- | --- | --- |
| 1 | Wall-associated kinases (*WAK/WAKL*) | involves in signal transduction between the cell wall and the cytoplasm in plants | Li et al. (2022) |
| 2 | protein MALE DISCOVERER 2 | recognizes the conserved signature motif of serine-rich endogenous peptides (SCOOPs) from microbes and elicits various immune responses | Hou et al. (2021) |
| 3 | SNF1-related protein kinase (*SnRK3*) | plays a role in the energy signaling and metabolism acting as metabolite sensors | Halford & Hey 2009; Jossier et al. (2009); Crozet et al. (2014) |
| 4 | Receptor-like protein kinase (*RLCK-VI*) | participates in pathogen-associated molecular pattern (PAMP)- pattern-triggered immunity (PTI) and effector-triggered immunity (ETI) | Lu et al. (2010); Zhang et al. (2010) |
| 5 | Casein kinase 1-like protein (*CKL*) | regulators of signal transduction pathways in most eukaryotic cell types | Wang et al. (2011) |
| 6 | Leucine-rich repeat receptor-like protein kinases (*LRR-VIII-2*) | contributes to plant defense against bacterial and fungal pathogen infection | Thapa et al. (2018) |

### Multiple sequence alignment and phylogenetic construction

To identify genetic variants such as single nucleotide polymorphisms (SNPs), simple sequence repeats (SSRs), and insertions and deletions (InDels) in partial sequences of *RLK* genes, multiple sequence alignments were generated per target gene using the ClustalW algorithm in Molecular Evolutionary Genetics Analysis (MEGA11) software (Tamura et al. 2021). Phylogenetic trees were also constructed using the maximum likelihood statistical method with the best DNA/Protein models (ML) (Kalyaanamoorthy et al. 2017), generated based on the lowest Bayesian Information Criterion (BIC) score, and evaluated with 1000 bootstrap replications in MEGA11.

### DNA marker development

To detect banana *RLK* genic variants across various *Musa* accessions, nine sets of gene-specific markers were developed using three marker systems—simple sequence repeat (SSR), allele-specific PCR (AS-PCR), and tetra primer amplification refractory mutation system (T-ARMS) PCR—using Geneious software (Kearse et al. 2012). Primer specificity and thermodynamics were further checked using Primer-BLAST (https://www.ncbi.nlm.nih.gov/tools/primer-blast) software and IDT OligoAnalyzer™ Tool (Integrated DNA Technologies, Inc., Coralville, Iowa; http://sg.idtdna.com/calc/analyzer).

#### Simple sequence repeat (SSR)

One SSR marker flanking the (TTC)_14_ motif of *RLCK-VI* was designed using the selected sequenced banana accessions, with an expected amplicon size of around 250 bp. Accessions with varying repeat unit number are expected to differ from the expected amplicon size. PCR products were resolved with electrophoresis using 8% non-denaturing polyacrylamide gel, and *Musa* individual samples were scored manually, with fragment lengths estimated using ImageJ software (Schneider et al. 2012).

#### Allele-specific PCR (AS-PCR)

Allele-specific PCR, also referred as PCR allele-specific amplification (PASA) or amplification refractory mutation system PCR (ARMS-PCR) markers (Newton et al. 1989; Little, 2001; Bundock et al. 2005), was one of the marker systems utilized for SNP detection. This marker system is comprised of one common primer pair [outer forward (OF) and outer reverse (OR)] and two inner forward (IF) [or inner reverse (IR)] allele-specific primers overlapping the SNP alleles with their 3’ terminal base, requiring an exact or perfect complementation in the last five bases of the template and allele-specific primer for successful primer extension (Gaudet et al. 2009). A deliberate mismatch at the penultimate (or second to the SNP allele) position was introduced in the inner primers following the required destabilization strength of nucleotide mismatch pairings recommended by Little (2001) and Bui and Lui (2009) to create a double mismatch to further increase allele specificity and discrimination power of the markers (Lefever et al. 2019). The rule of thumb is that if the identified SNP mismatch has a weak destabilizing strength, then a strong destabilizing deliberate mismatch at the penultimate position of the SNP should be introduced in the primer, and vice versa. Conversely, if there is a medium destabilizing SNP mismatch, a weak or medium destabilizing mismatch should be introduced at the penultimate position of the primer’s 3’ terminus. As a result, the correct mismatch base was introduced in each inner primer. The assay is composed of two PCR rounds: (1) amplification of control fragment (longer fragment) containing the region of SNP site using outer primers and the amplicon product was subsequently used as a template (2 uL) in the second PCR round, and (2) simultaneous (parallel) reaction assay of SNP alleles using AS primers (inner primers) paired with the opposite outer primer (Darawi et al. 2013). The PCR component concentrations and profile are the same in the aforementioned PCR methods and amplicons were resolved in 8% non-denaturing polyacrylamide gel. The banding patterns of alleles were assessed through manual scoring.

#### Tetra primer amplification refractory mutation system (T-ARMS) PCR

T-ARMS PCR markers, another SNP detection assay, were designed using four primers consisting of one outer primer pair (OF/OR) and two inner primers (IF/IR), asymmetrically designed with respect to the outer primers and coamplifying target SNP alleles in a single PCR reaction. Consequently, outer primers are intended to amplify the control fragment to ensure optimal PCR conditions. Since this marker system is a multiplex PCR, there is a potential for primer interactions and competition in PCR components, which could affect marker’s efficiency and specificity and necessitate multiple rounds of optimization. To identify the most efficient PCR condition, several concentrations of various PCR components (e.g., inner primers, MgCl_2_, *Taq* polymerase, etc.) and different annealing temperatures were tested. The optimized PCR reaction had a total volume of 10 uL consisting of 1X PCR buffer (10 mM Tris pH 9.1 at 20°C, 50 mM KCl, 0.01% Triton™ X-100, Vivantis Technologies, Malaysia), 2.5 mM MgCl_2_, 0.2 mM dNTPs (Promega Corporation, Madison, Wisconsin, USA), 1:5:10 ratio for outer/inner forward/inner reverse primer concentrations, 0.5 U recombinant Taq DNA polymerase (Vivantis Technologies, Malaysia), and 10 ng DNA. PCR products were resolved using gel electrophoresis at 100 V for 40 min on 1.5% agarose gel. Banding patterns from gels were scored codominantly.

### Validation of *RLK*-linked DNA markers

The efficiency and polymorphism of nine gene-specific markers were tested and validated across 117 *Musa* accessions selected from NPGRL-IPB *ex situ* germplasm collection. This comprehensive set included various accessions, such as 29 wild *M. balbisiana* accessions, one wild *M. acuminata* (‘Agutay’), ten diploids (AA, BB), 52 triploid bananas (AAA, AAB, ABB, BBB), eight tetraploid accessions (AAAA, AAAB, ABBB), seven other *Musa* species (including ornamental bananas, putative interspecific banana accession ‘Cantong’, and ‘Abaca’), and 10 banana genotypes with unknown genomic constitution (see Supplementary Table S2). The genotypic profiles of the banana population were successfully generated through allele and repeat unit number scoring using positive control samples (banana accessions with consensus nucleotide sequences). Polymorphism indices of DNA markers such as allele and genotype frequencies, number of alleles per locus (Na), number of effective alleles (Ne), observed (Ho) and expected (He) heterozygosity, and Wright’s fixation index [inbreeding coefficient (FIS) only] were analyzed and calculated using GenAlEx ver. 6.51b2 software (Peakall and Smouse 2012) with manual computations. Polymorphic information content (PIC) was computed using an online tool at https://www.gene-calc.pl/pic.

### Genetic diversity analysis

Using the developed polymorphic SNP markers, a dendrogram was constructed using DARwin (Dissimilarity Analysis and Representation for Windows) 6 software (Perrier and Jaquemoud-Collet 2006) with default dissimilarity index of allelic data by Unweighted Neighbour-Joining (UNJ) method with 1,000 bootstrap iterations. The graphic representation was formatted using FigTree v1.4.4 (Rambaut et al. 2019) software.

### Assessment of the functional impact of *RLK* variants

To initially assess the roles of *RLK* gene variation in disease resistance, genomic localization of SNPs and SSR, substitution type, and amino acid change were determined by checking the correct reading frame of nucleotide sequences per gene using the ExPASy translate tool (https://web.expasy.org/translate/) for translation to polypeptide sequence.

### Marker-trait association analysis

A sampling strategy was implemented to collect a diverse representative of 117 *Musa* genotypes (see Supplementary Table S2) for association analysis with respect to BBTV incidence data. In a natural field setting characterized by BBTV prevalence, three to five banana suckers were collected for each individual accession. Leaf samples from each banana genotype were then pooled, and an established DNA isolation method was undertaken to extract high-quality genomic DNA for each genotype using Doyle and Doyle (1990) method with modifications. The initial phenotypic assessment of BBTV presence or absence was executed via a quantitative polymerase chain reaction (qPCR) detection assay for accurate detection of BBTV infection status of the samples as described by Mendoza et al. (under review). To prepare the data suitable for statistical analysis, the amplification curves from the qPCR assay were transformed into binary data. Single marker analysis, using Generalized Linear Model (GLM) function incorporated into the RStudio platform (RStudio Team 2020), was performed to check association between polymorphic SNP markers and BBTV presence in natural field conditions.

## RESULTS

### Banana *RLK* variants

To investigate *RLK* gene sequence diversity and polymorphisms across banana accessions, sequencing primers per target *RLK* gene were developed (see Supplementary Table S1), and their respective gene locations are presented in Fig. 1a. These sequencing primers were effectively optimized and amplified the target gene regions (Fig. 1b), yielding distinct and high-quality amplicons required for outsourced partial sequencing. Through standard sequence quality control, high-quality consensus sequences for each *RLK* gene were generated for subsequent analyses such as multiple sequence alignment and diversity analysis.

**Fig. 1.**
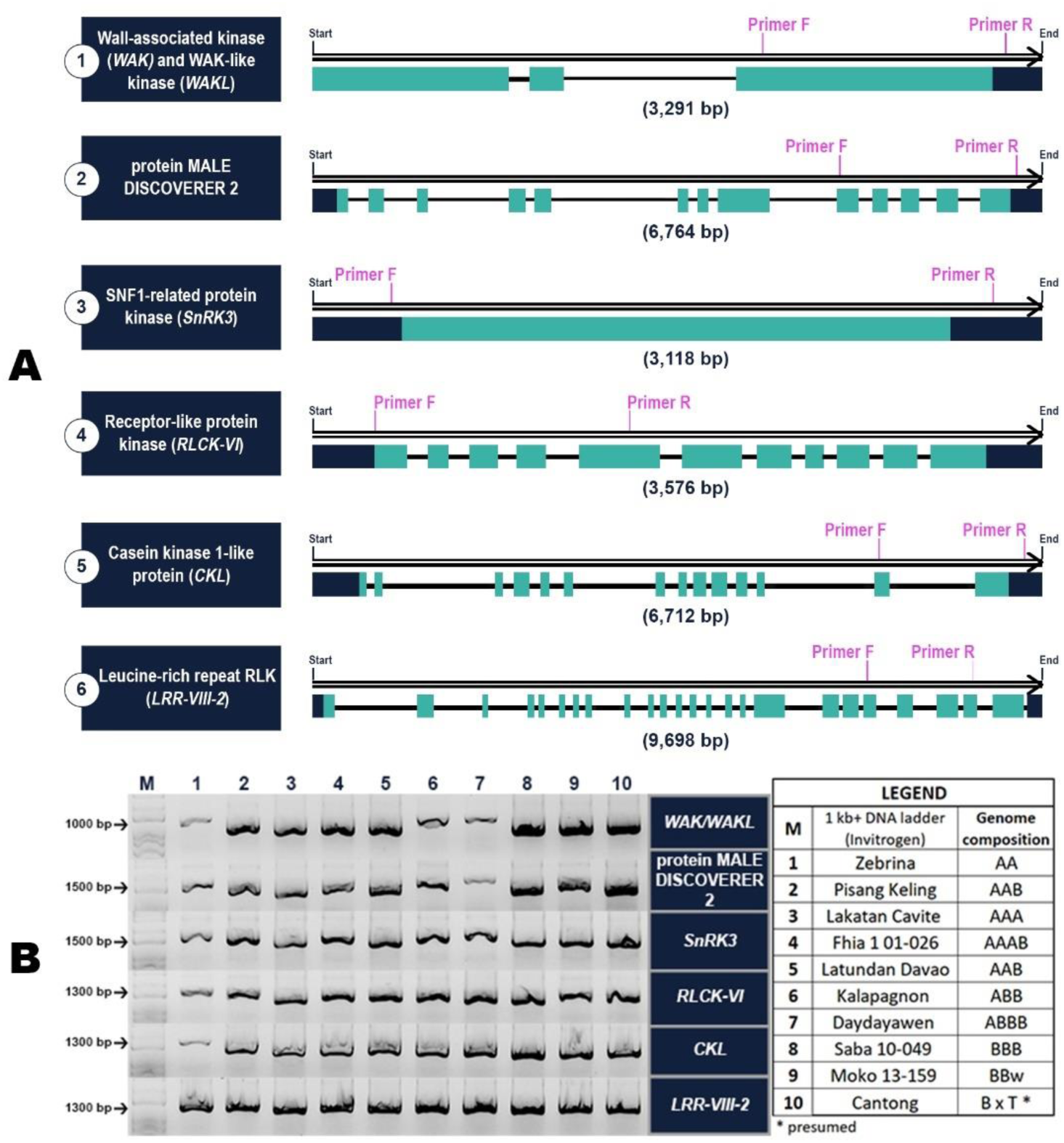
Sequencing primers for each *RLK* genes: a) gene structure of *RLK* genes retrieved from the Banana Genome Hub (https://banana-genome-hub.southgreen.fr/) and location of sequencing primers; b) Amplified PCR products of six *RLK* genes in 10 banana accessions for partial gene sequencing. Gene features color legends: dark blue – untranslated regions (UTRs), turquoise – exons, and black – introns.

Multiple sequence alignment analysis of each *RLK* gene consensus sequence derived from 10 *Musa* accessions revealed variable sites such as simple sequence repeat (SSR) and single nucleotide polymorphisms (SNPs), both are abundant in plant genomes and provide high degrees of polymorphisms (Choudhury et al. 2023). The summary of genic variants found in six banana *RLKs* is presented in (Supplementary Table S3). The majority of the generated *RLK* gene partial sequences contain single nucleotide polymorphisms (SNPs) because they are the most common and abundant form of genetic variation in plants (Liao and Lee 2010; Mammadov et al. 2012). A substantial number of SNPs were also observed in the introns compared to exons, as this is predominantly present in non-coding than in coding regions of plant genomes (Manrique-Carpintero et al. 2013; Zhang et al. 2018). Meanwhile, while SSRs are also generally abundant in plant genomes, only one repeat motif was found in one of the *RLK* gene sequences and was located in the intronic region of *RLCK-VI*, as expected given the relatively lower abundance of SSRs in the coding regions due to their high mutation rate that could affect gene expression or functionality of proteins (Vieira et al. 2016). The identification of genetic variations in BBTV-responsive *RLK* genes in selected banana accessions will lay the groundwork for DNA marker development.

### Marker development for *RLK* genes

Using the identified *RLK* genic variants (SSR and SNPs), a total of nine sets of gene-specific markers were developed using three distinct marker systems: SSR, AS-PCR, and T-ARMS PCR, enabling detection of *RLK* genetic variations across various *Musa* accessions. Table 3 outlines the marker systems and primer sequences employed for each target gene while Fig. 2 depicts the representative schematic diagrams of marker design for each marker system and the resulting electrophoretograms.

**Fig. 2.**
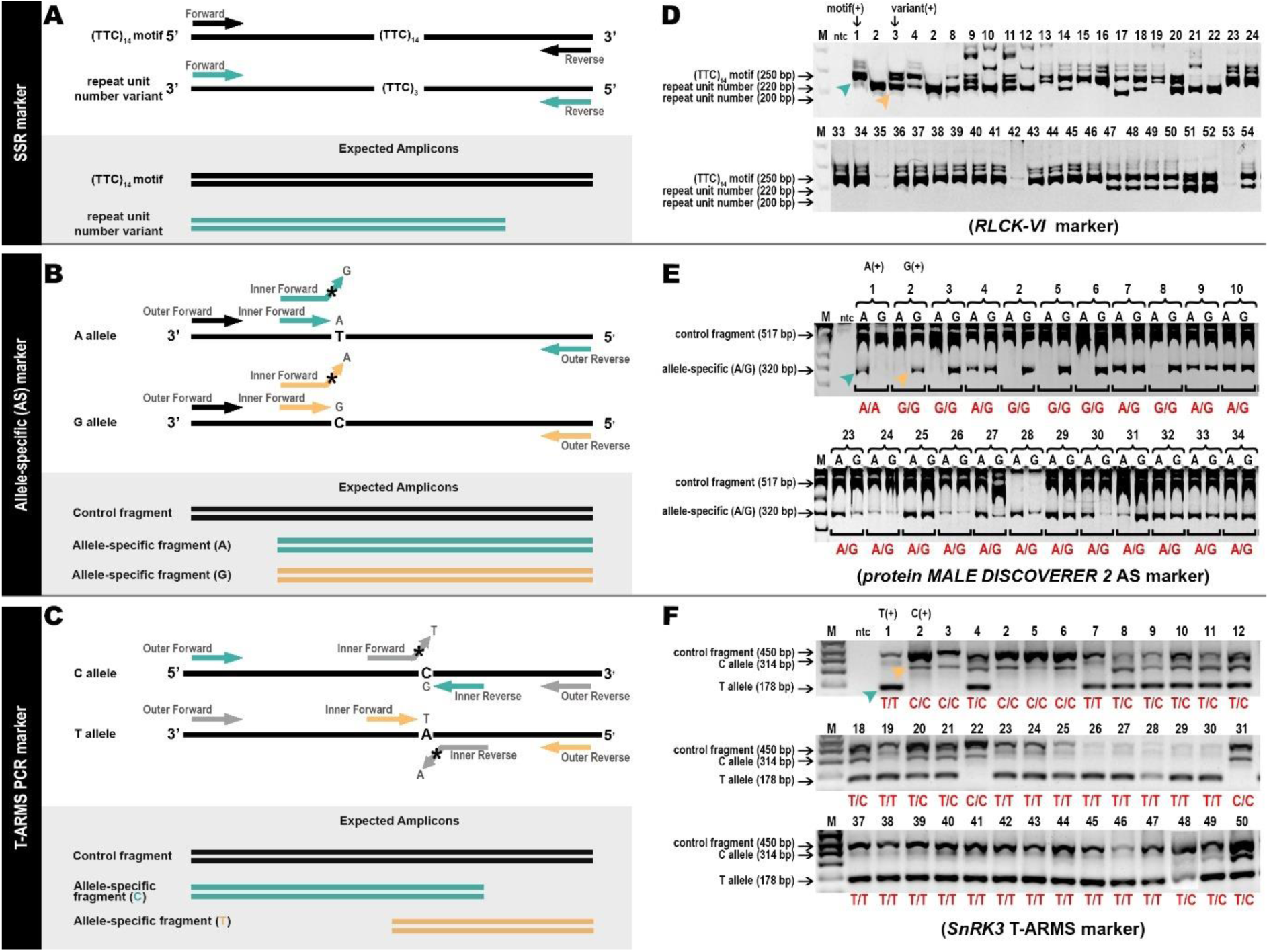
Representative schematic representation of each marker system and resulting electrophoretograms for each marker type. Figure 2a-c are sample marker designs for SSR marker (a), AS-PCR marker (b), and T-ARMS PCR marker (c) and their expected amplicon sizes. Asterisk represents the deliberate mismatch base introduced at the penultimate position of SNP site. Figure 2d-f are representative electrophoretograms of selected *RLKs* for each marker system: resulting electrophoretogram of SSR marker for *RLCK-VI* (d), AS-PCR marker for protein MALE DISCOVERER 2 (e), and T-ARMS PCR marker for *SnRK3* (f). Lane M stands for 1 kb Plus DNA ladder (Invitrogen)(1.5% agarose gel) (Fig 2f)/DNA molecular weight marker VIII (8% denaturing polyacrylamide gel) (Fig 2d-e), lane 1 refers to the genotype of BBTV-resistant wild *M. balbisiana*, lane 2 refers to the genotype of BBTV-susceptible banana cv. ‘Lakatan’, turquoise and tan arrows refer to positive control samples’ allele (based on nucleotide sequences), and banana accessions corresponding to the lane numbers/sample code are presented in Supplementary Table S2.

**Table 3.**
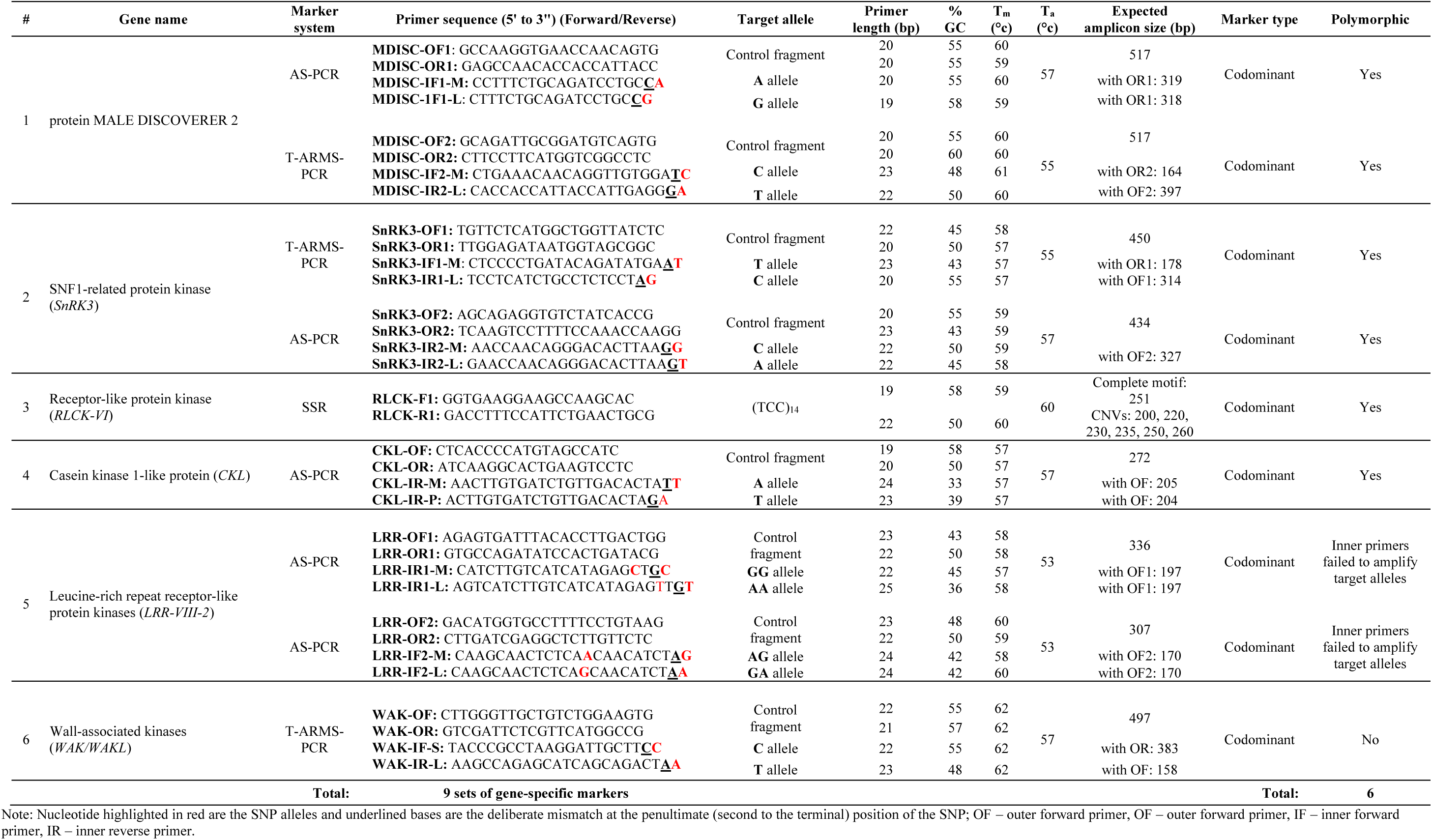
Gene-specific markers developed from partial gene sequences of six *RLK* genes in representative bananas.

One (*RLCK-VI*) of the six *RLK* genes from representative bananas revealed an intronic SSR region with (TTC)_14_ repeat motif and repeat variation across different banana genotypes. As depicted in Fig. 2a, SSR marker was designed flanking the repeat motif, enabling the detection of repeat unit number variation banana genotypes. On the other hand, AS-PCR were designed to be highly specific to the target SNP site through addition of deliberate mismatch to the primer (represented by an asterisk in Figure 2b), resulting in the presence or absence of the allele of interest. Moreover, T-ARMS PCR simultaneously detects target SNP alleles in a single PCR reaction through asymmetric design of the allele-specific primers, obtaining amplicon size variation, as shown in Figure 2c. Following marker design, all of these were efficiently optimized for utilization in banana genotyping.

### Validation of *RLK* DNA markers

To validate the effectiveness of the established markers for genotyping assay, markers were tested across 117 *Musa* genotypes from NPGRL-IPB *ex situ* germplasm collection for their capability to detect genetic variations or polymorphisms of *RLK* genes. Six (one SSR, three AS-PCR, and two T-ARMS PCR markers) out of the nine developed DNA markers were polymorphic and can discriminate various *Musa* accessions, whereas the remaining three markers were monomorphic. Fig. 2d-f shows the representative electrophoretograms of the polymorphic markers.

The SSR marker efficiently identified different alleles and variability in repeat numbers within the *RLCK-VI* repeat motif region, yielding unique bands for each identified allele in banana accessions (Fig. 2d). Detection of the expected amplicon sizes based on the repeat motif’s number variations in the positive control samples, BBTV-resistant wild *M. balbisiana* accession ‘Moko 13-159’ (250 bp, turquoise arrow) and ornamental banana ‘Zebrina’ (AA) (220 bp, tan arrow), facilitated successful genotyping scoring of the sampled population. Despite the fact that BBTV-susceptible *M. acuminata* cv. ‘Lakatan’ (AAA) was not included in the multiple sequence alignment of *RLCK-VI* sequences due to noisy and low-quality sequence reads, the marker still obtained the expected amplicon size (220 bp) as shown in Fig. 2d lane 2, which is similar to the repeat unit number of ‘Zebrina’ (AA), as both have the same genome composition differing only on the ploidy level. Other repeat unit number variations were also revealed in various banana accessions, demonstrating effective amplification across diverse banana accessions.

Furthermore, three of the five established AS-PCR markers—protein MALE DISCOVERER 2, *SnRK3*, and *CKL* markers—with target SNPs A796G, C724A, and A982T, respectively, were also identified to be polymorphic across the *Musa* population, demonstrating high specificity of the markers to the target SNP alleles. Codominant scoring of the *Musa* genotypes was effectively facilitated by successful detection of the predicted allele bands of the positive control genotypes [‘Moko’ (BBTV-resistant) and ‘Lakatan’ (BBTV-susceptible)], as depicted in Fig. 2e (AS-PCR marker for protein MALE DISCOVERER 2).

Likewise, two of the three T-ARMS markers, targeting the allele variations of protein MALE DISCOVERER 2 (SNP C1068T) and *SnRK3* (SNP T943C) genes, successfully differentiated various *Musa* genotypes. Polymorphic markers displayed the expected genotypes of the positive controls demonstrating complete concordance to the generated SNP sequences, enabling SNP genotyping of banana accessions via co-dominant scoring (Fig. 2f). Unfortunately, the T-ARMS PCR marker for the *WAK/WAKL* gene displayed a monomorphic banding pattern. It is suggested that the primer be redesigned or other SNP detection methods (e.g., KASP genotyping, TaqMan probes) be implemented. The complete genotypic profiles of the 117 *Musa* accessions using six polymorphic markers are presented in Supplementary Figs. S1-6.

Overall, the established polymorphic markers (SSR, AS-PCR and, T-ARMS PCR) exhibited high discriminatory power across the banana population and provided a robust and cost-effective technique to identify genic variations on the targeted genes.

### Polymorphism indices of six *RLK* DNA markers

The polymorphism information of six *RLK* DNA markers was generated and presented in Table 4. Due to the bi-allelic nature of SNP markers, the total number of alleles (Na) and effective number of alleles (Ne) for all five allele-specific markers are equal to two. On the other hand, SSR marker for *RLCK-VI* contains a total of six alleles (Na); however, the effective number of alleles (Ne) computed was only two alleles. Although SSRs are multi-allelic in nature, the detected number of effective alleles (Ne=2.14) in 117 *Musa* accessions might be due to the presence of A- and B-genomes, with most alleles scoring codominantly, whereas the remaining four alleles account for the rare and private alleles coming from other *Musa* spp. like ornamental bananas and abaca. While tri-allelic accessions are relatively rare, they are particularly observed in triploids and tetraploids with a prominent fraction of the A genome (AAA, AAB, AAAA, and AAAB). All accessions of wild *M. balbisiana* showed homozygous alleles for the (TTC)_14_ motif while wild *M. acuminata* (‘Agutay’) and other accessions of *M. acuminata* (i.e., ‘Lakatan’, ‘Mapilak’, ‘AACV Rose’, etc.) displayed homozygous alleles for its repeat unit number (shorter fragment), and accessions with A- and B-genome combinations revealed heterozygous genotype. In addition, the major and minor allele frequencies for the SSR marker are 0.510 and 0.006, respectively (see Supplementary Tables S4 and S5). The major allele (E allele) was present in all wild *M. balbisiana* accessions as well as hybrid bananas (triploids and tetraploids) containing B genomes. Minor or rare alleles (C and F alleles) were discovered in ornamental bananas (‘Ornata’ accessions 16**-**017 and 98-156). The next lowest minor allele frequency (MAF) was found in banana accessions ‘Abaca’, ‘Cantong’, ‘Pakil’, and ‘Paquel’ with a frequency value equal to 0.026. This may indicate that there is a possibility that ‘Cantong’, ‘Pakil’ and ‘Paquel’ are more related to abaca (*Musa textilis*) as they shared unique alleles. This further reinforces the presumption that ‘Cantong’ is a natural hybrid between *M. balbisiana* and *M. textilis*. Nonetheless, additional investigation is needed to definitively conclude the notion.

**Table 4.**
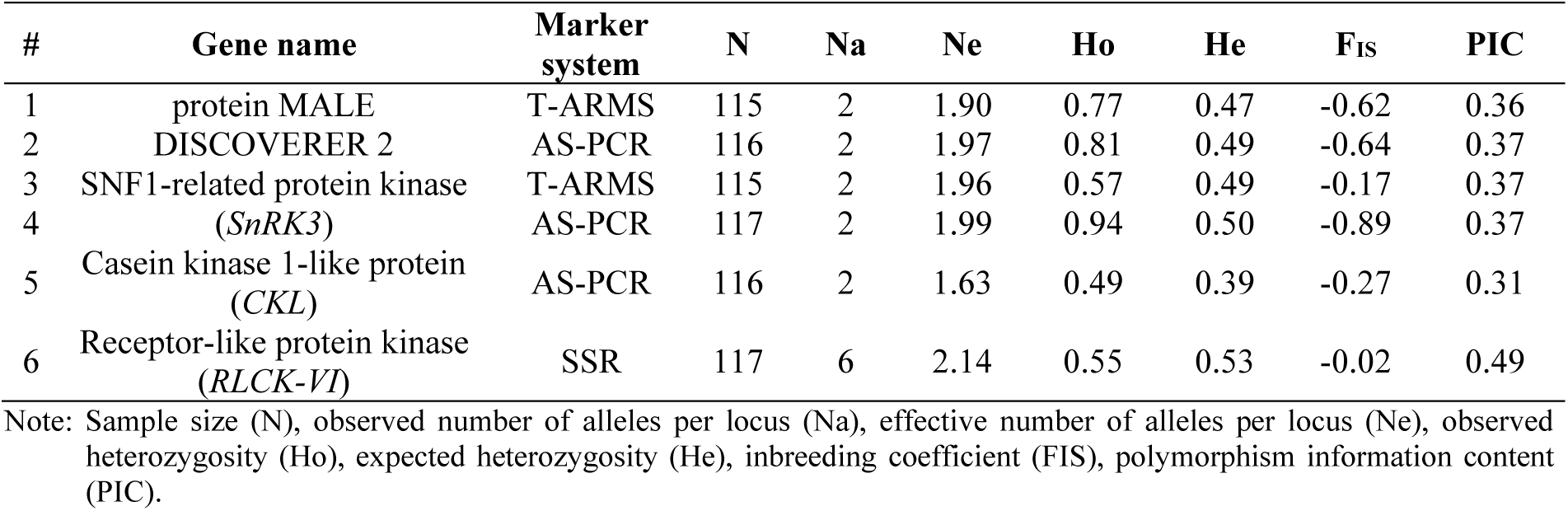
Polymorphism indices summary of six *RLK* gene-specific markers in 117 *Musa* accessions.

Furthermore, due to the high complexity of the *Musa* genome composition and ploidy levels, the observed heterozygosity (Ho) for all six DNA markers ranges from 0.49 to 0.94, which is greater than the expected heterozygosity (He: 0.39-0.53). As a result, the population’s fixation index or inbreeding coefficient (FIS) was negative for all markers indicating high or excess heterozygosity (Meisner and Albrechtsen 2019) in the sampled natural population which is consistent with the findings of Gardoce et al. (2023). The polymorphism information content (PIC) for all markers ranges from 0.31 (*CKL* AS-PCR marker) to 0.49 (*RLCK-VI* SSR marker). The average PIC for the five SNP markers (AS-PCR and T-ARMS PCR) and one SSR marker is approximately 0.36 and 0.49, respectively, both of which are moderately informative and are useful for discriminating banana genotypes.

### Genetic diversity

Using the polymorphic SNP markers, the constructed dendrogram identified four major clusters of the *Musa* population in correspondence to their genomic compositions or predetermined groups (Fig. 3). The A-genome accessions with different ploidy levels (i.e., diploids, triploids, and tetraploids), along with few inter-specific *Musa* hybrid accessions comprising A- and B-genomes, formed a major cluster (I) consisting of four subgroups. Within this major cluster, *M. acuminata* accessions ‘Agutay’ (AAw), ‘Lakatan’ (AAA), and other *Musa* ssp. ‘Laterita’ formed a unique group, clustering out from the diploids (AA) such as ‘Zebrina’, ‘AACV Rose 13-16’, ‘Cuarenta Diaz’, and ‘Eda-an’. Intriguingly, ‘Mapilak’ (AAA) which is a mutant of *M. acuminata* cv. ‘Lakatan’ (AAA), also clustered together with the diploids (AA) and dissociated from the ‘Lakatan’ variety, which may suggest that some of the targeted alleles of the developed markers here were mutated in one of the genome copies in the BBTV-resistant banana mutant accession ‘Mapilak’, presenting a potential research direction for the study of the functional effect of mutation in *RLK* genes. In addition, most of the triploids and tetraploids with purely A-genome were sub-grouped together, with some admixture of other *Musa* spp. like ornamental bananas, and clearly separated from the diploid accessions. A few triploids and tetraploids comprising a combination of A and B genomes also established a subgroup in proximity of the *M. acuminata* accessions.

**Fig. 3.**
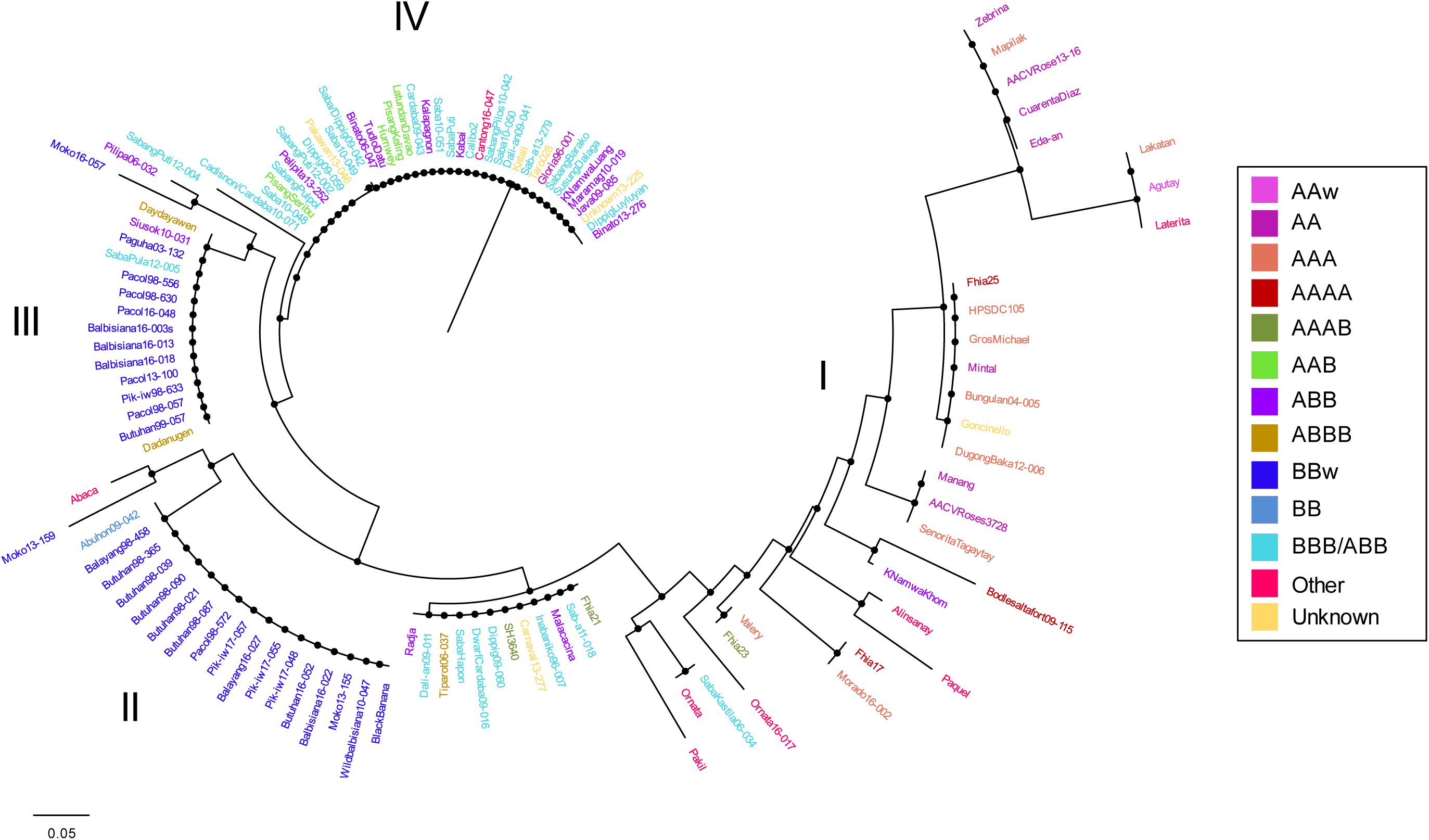
Dendrogram generated by Unweighted Neighbor-Joining (UNJ) method with 1,000 bootstrap iterations using dissimilarity index allelic data of five SNP markers, forming four major clusters (I, II, III, and IV) revealing the genetic relationship of the 117 *Musa* spp. accessions from IPB-NPGRL *ex situ* germplasm collection.

Moreover, wild *M. balbisiana* (BBw) accessions formed two major clusters (II & III) and separated completely from the *M. acuminata* accessions, with the inclusion of some triploid and tetraploid accessions having a significant proportion of the B-genome (i.e., ABB and ABBB). The formation of two major groups within the wild *M. balbisiana* accessions is due to its intra-population genetic diversity. Interestingly, one accession of *M. textilis* (‘Abaca’) formed a group with the wild *M. balbisiana* accessions coinciding with the results of Gardoce et al. (2023). Nonetheless, hybrid bananas with varying ploidy levels also established a significant cluster (IV) due to heterozygosity scores, regardless of their A or B-genome dosage, in the target alleles. These findings clearly demonstrated the ability of these markers to discriminate major *Musa* genomic groups in the population and successfully revealed the genetic relationships of the banana genotypes.

### *Ex situ* germplasm management

The established molecular markers here can also be used as genome specific markers to identify B-containing genome from A-containing genome banana accessions, emphasizing its utility in *ex situ* banana germplasm management. Two DNA markers, namely *RLCK-VI* SSR and *SnRK3* T-ARMS markers, effectively discriminated wild *M. balbisiana* accessions (B-genome) from other banana accessions such as *M. acuminata* (A-genome) and hybrids (triploids and tetraploids). Further, three allele-specific DNA markers (*protein MALE DISCOVERER 2* AS-PCR and T-ARMS PCR marker, and *SnRK3* AS-PCR marker) can distinguish banana accessions containing B-genome. Although there are some banana accessions that show unexpected allele-scoring patterns which require further research and re-evaluation of genomic compositions and passport data, these DNA markers could be used to identify and further validate banana accessions in the field, as well as other newly acquired germplasm accessions.

### Functional significance of *RLK* gene variations

The functional importance of *RLK* variations in connection to BBTD resistance was assessed preliminary by identifying the type of base substitution, gene location (e.g., exon, intron, UTRs, etc.), and degree of variant impact in gene expression, mRNA stability, and protein function influencing population phenotypic variations (Gu et al. 2015). Two types of DNA variations identified in the *RLK* gene fragments are SSRs and SNPs which can be found in the coding and non-coding regions of the genome and may have further significant effects on protein functionality depending on the impact level of the genic variants. Table 5 presents the summary of SSR and SNP location, substitution type, and the corresponding amino acid change identified in four *RLKs* with polymorphic DNA markers. Repeat motif of the SSR was identified in the intron region of the *RLCK-VI* while majority of the SNP markers were targeted in the exonic region of the genes. Three of the five SNPs found in the *RLK* genes (protein MALE DISCOVERER 2, *SnRK3*, and *CKL*) showed non-synonymous substitution, displaying a change of amino acid from serine (S) to glycine (G), glutamine (Q) to histidine (H), and aspartic acid (D) to glutamic acid (E), while the remaining two (protein MALE DISCOVERER 2 T-ARMS PCR and *SnRK3* AS-PCR markers) showed synonymous substitution or silent mutation.

**Table 5.**
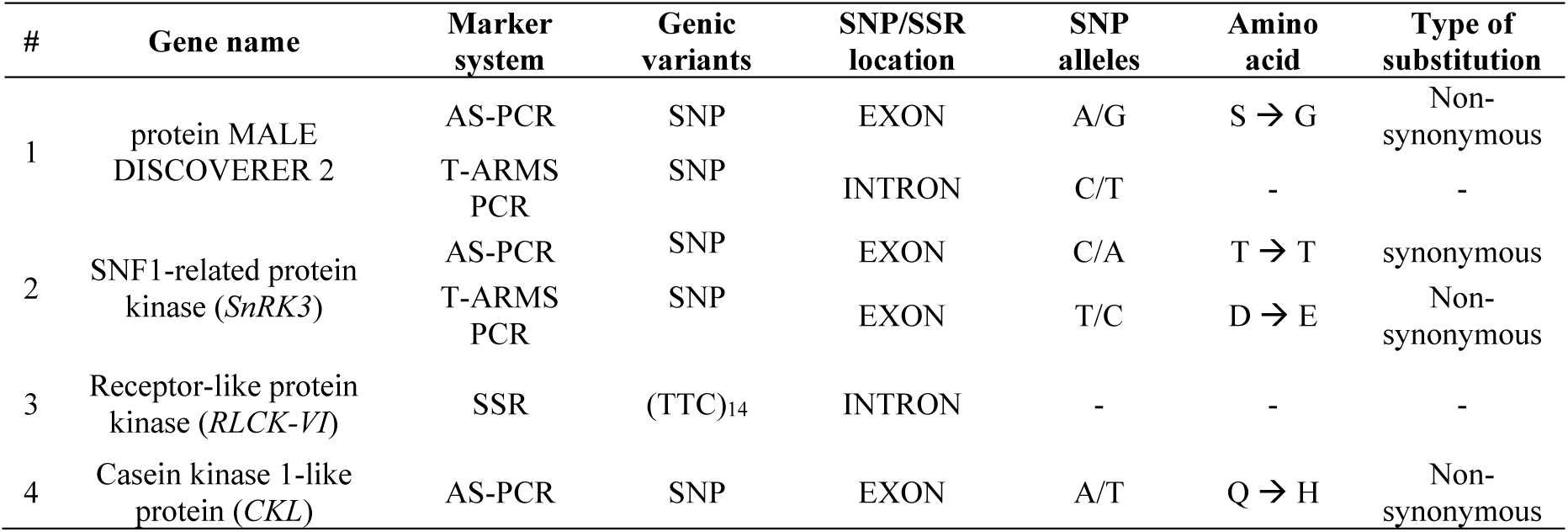
Summary list of SSR and SNP genic variants of *RLK* genes in the *Musa* genome including their location, alleles, and types of substitution identified in this study.

### Marker-trait association analysis

To provide initial insights into the potential association of polymorphic SNP markers with BBTV resistance, single marker analysis was performed in the genotype and phenotypic data of the selected *Musa* accessions. Phenotypic data of the *Musa* accessions using natural field infections (presence or absence of BBTV using qPCR detection; Fig. 4) were initially gathered, allowing initial assessment of individual DNA markers and their potential association with BBTV resistance of bananas. Table 6 showed that only one of the five SNP markers (*SnRK3* T-ARMS PCR marker) revealed a statistically significant association to BBTV incidence, with a p-value of 0.0162, at 0.05 significance level.

**Fig. 4.**
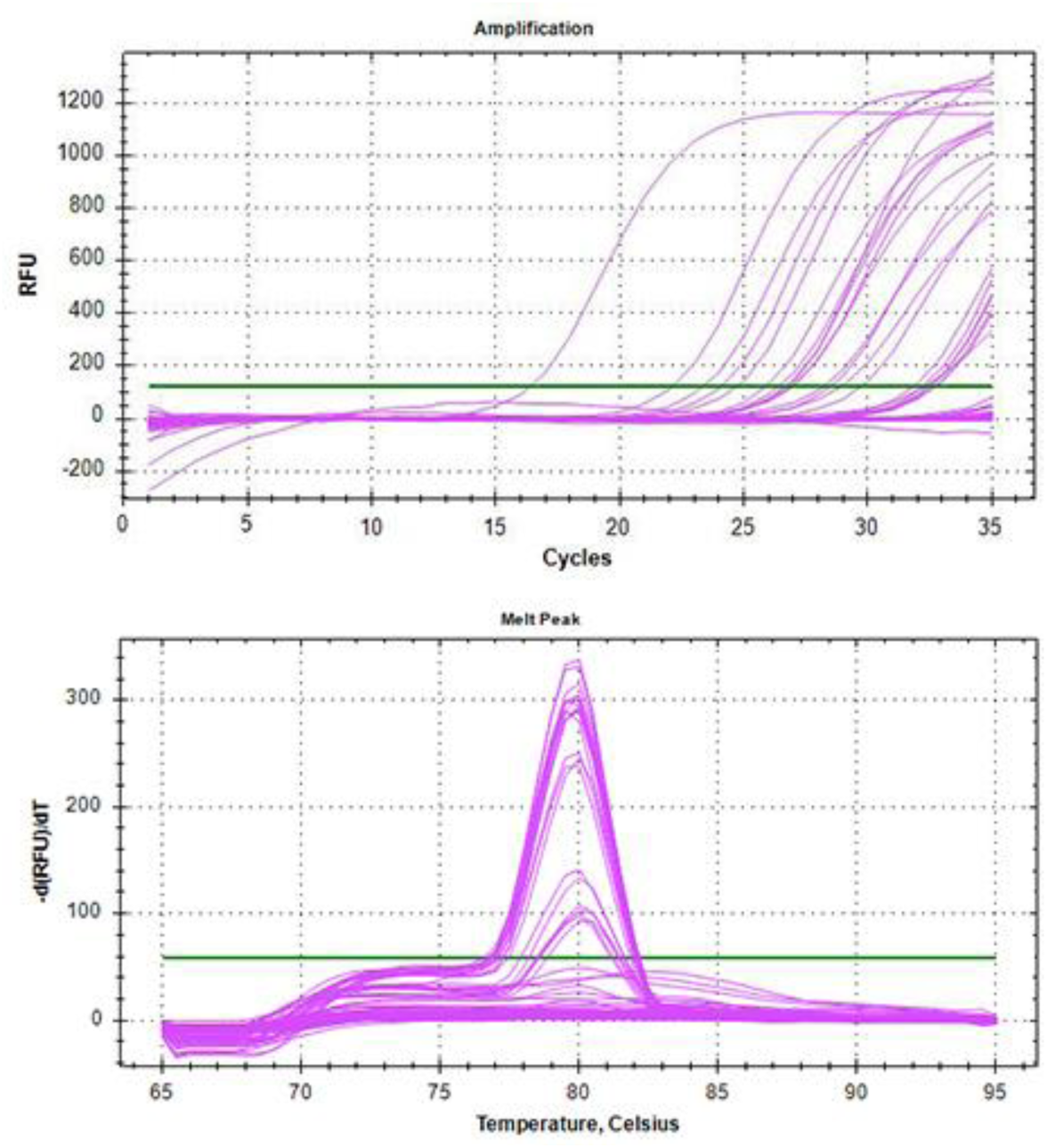
qPCR amplification curves, targeting *DNA-R* region of the *banana bunchy top virus*, across 98 banana accessions. The curves illustrate the exponential increase in fluorescence signal with each cycle, indicating the presence of viral genetic component.

**Table 6.** Single marker analysis of SNP markers and initial phenotypic data obtained from banana bunchy top virus detection in naturally infested field conditions.

| Entry # | Marker Name | Standard error | Z value | Pr(>\|z\|) |
| --- | --- | --- | --- | --- |
| 1 | MDISCTET | 0.63367 | -0.033 | 0.9737 |
| 2 | MDISCAS | 0.55991 | -0.098 | 0.9216 |
| 3 | SNRK3TET | 0.47811 | 2.404 | 0.0162 * |
| 4 | SNRK3AS | 1.25906 | -0.773 | 0.4393 |
| 5 | CKLAS | 0.47715 | 0.359 | 0.7197 |

Significance level codes: 0 '\*\*\*\*' 0.001 '\*\*\*' 0.01 '\*\*' 0.05 '\*'

## DISCUSSION

Through extensive transcriptome analysis, the recent findings of Lantican et al. (2023) have expanded our understanding on the molecular mechanisms underpinning BBTV resistance in bananas, paving the way for targeted genetic improvement to expedite the development of BBTV-resistant banana cultivars. To continue this initiative, we further investigated gene sequence diversity and polymorphisms of six *RLK* genes, the first line of plant defense against pathogens, across banana accessions at the NPGRL-IPB’s banana *ex situ* germplasm collection. We have identified two genic variants of banana *RLKs,* namely SNPs and SSR, which have been the basis of molecular marker development, enabling successful genotyping of bananas.

Molecular markers, such as SSR and SNPs, are widely used in various genotyping applications as these are excellent tools in genetic diversity studies and linkage analyses due to their inherent versatility, ease of assay, and abundance in plant genomes (Jiang 2013; Marwal et al. 2014). Here, one SSR marker has been developed targeting the repeat unit number variation of the *RLCK-VI* gene and successfully identified distinct alleles across various *Musa* genotypes. On the other hand, a simple and cost-effective PCR and gel-based marker systems such as allele-specific PCR (AS-PCR) and tetra primer amplification refractory mutation system PCR (T-ARMS PCR) markers were used as an alternative approach to the current SNP detection technologies such as genotyping-by-sequencing (GBS) approach, SNP microarrays, and PCR-based fluorescently labeled high-throughput methods like fluorescent probes and kompetitive allele-specific PCR (KASP) (Thomson 2014; Semagn et al. 2014; Matsuda 2017), as these are quite costly for routine use and require specialized equipment.

AS-PCR and T-ARMS makers identify SNPs by selective amplification of target SNP alleles, resulting in the presence or absence of amplicon products. Three of the five developed AS-PCR markers were polymorphic across the banana population, demonstrating high specificity to the target SNP alleles. Meanwhile, amplification failure of the remaining two AS-PCR markers (*LRR-VIII-2* AS-PCR markers), which was designed with an additional adjacent SNPs in the last five bases at the 3’ end of the target DNA template (highlighted bases in Table 4), provided us with significant insights into its implication on amplification success. The presence of two adjacent SNPs near the 3’ end when coupled with deliberate mismatch next to the 3’ SNP termini, resulted to ineffective amplification of the target SNP allele due to an increase in destabilization strength of the mismatch base pairings affecting primer annealing and extension. Hence, this factor should be considered when designing allele-specific DNA markers and we recommend targeting only single SNP site at a time.

Likewise, T-ARMS PCR allows SNP detection similar to AS-PCR but utilizes a different and improved approach in discriminating specific target alleles. Unlike AS-PCR, this method simultaneously detects the target SNP alleles in a single PCR reaction, instead of several or parallel PCR reactions. However, because it uses four primers in a single reaction, this marker system necessitates rigorous optimization process such as primer concentration counterbalancing, which we achieved at 1:5:10 ratio for outer/inner forward/inner reverse primers, similar to the recommended ratio of Ye et al. (2001) that is 1:10 (outer/inner primer), with slight difference on inner primer concentration due to preferential amplification between the shorter and longer allele-specific fragments. Despite the difficulty in optimization, we successfully produced two polymorphic markers from three T-ARMS PCR markers developed, while one marker (*WAK*/*WAKL* T-ARMS PCR) displayed monomorphic banding pattern. One of the key causes for the failure of T-ARMS-PCR method to discriminate target SNPs is low specificity of the primers, increasing the chances of cross-reactivity with other alleles or non-target SNP sites. Nonetheless, both allele-specific markers (AS-PCR and T-ARMS PCR) successfully detected target SNP alleles in the banana population.

Although SSR markers provide high allelic diversity and co-dominant inheritance, SNP markers offer higher throughput and are suitable for large-scale genotyping assays (Wang et al. 2021; Wu et al. 2021). Both AS-PCR and T-ARMS PCR markers provide excellent specificity and sensitivity for SNP genotyping polyploid crops, but the use of T-ARMS PCR markers is more cost-effective and time-efficient, making it particularly attractive for large-scale genotyping studies. Overall, the established six polymorphic markers (one SSR, three AS-PCR, and two T-ARMS PCR) in this study exhibited sufficient discriminatory power across the tested banana population and successfully demonstrated their ability to distinguish major banana genomic groups, revealing the genetic diversity and relationships of *Musa* genotypes which are useful in germplasm management and downstream association studies.

Moreover, the impact level of the identified SNPs was determined and classified based on the annotation of Bohry and collaborators (2021) as low (e.g., synonymous variant or silent mutation), medium (non-synonymous mutation or missense mutations), and high (e.g., loss or gain of stop codon and frameshift variants) to further understand the functional significance of *RLK* variations in banana. Because only silent and missense mutations and none of the high impact variants that cause protein function loss were observed, SNPs identified in this study have low to moderate impact on *RLK* gene function which may still affect the efficiency of disease resistance function of these genes. These preliminary findings may help us better understand the impact of these mutations on the gene’s ability to combat banana bunchy top disease. However, it is vital to underline the importance of additional functional studies such as gene silencing, knockout, and overexpression (Bao et al. 2022) to further support and validate this inference.

Nevertheless, initial phenotypic data of the *Musa* accessions were obtained and used for marker-trait association analysis, providing information on the potential association of polymorphic SNP markers with BBTV resistance. One SNP marker targeting *SnRK3* gene was found to be associated with BBTV status or incidence in natural field. As previously summarized in Table 2, *SnRK3* is primarily involved in several metabolic pathways for energy conservation as coping mechanism for abiotic stresses, but it has also been linked to plant defense responses against pathogens (Wang et al. 2019), further implying that the same signaling pathways triggered in response to abiotic stresses could also potentially play a role in plant defenses against pathogens, leading to the activation of defense genes or synthesis of defensive proteins to counteract pathogen invasion. With this, the results obtained here provided initial insights on the possible role of *RLKs* (i.e., *SnRK3*) in the dynamics of disease transmission and host responses, as well as presented the potential genes associated with BBTD resistance.

The future direction of this study is to perform genome-wide association studies (GWAS) using high-density SNP markers and actual viral titers (based on qPCR Ct values) of banana accessions as phenotypic data, which will shed light on the intricate interplay between genetic variants and phenotypic traits associated with BBTV resistance.

## CONCLUSION

In conclusion, the markers developed in this study hold significant utility for *ex situ* banana germplasm maintenance and management, providing a valuable asset for banana genetic improvement initiatives, and for subsequent association studies focused on resistance to banana bunchy top disease, which will be later useful in marker-assisted plant breeding applications. Moreover, the assessment of *RLK* variations’ functional relevance could also offer insight on its disease resistance role against BBTV infection, although further exploration on the level of influence of these variants on protein structure, stability, and function is required. Nonetheless, initial marker association analysis provided us valuable insights on genes potentially linked with BBTD resistance, but further exploration on marker association using a data obtained from disease resistance screening and quantitative data from BBTV viral titer is recommended and will be the future direction of this study.

## ACKNOWLEDGMENTS

The authors would also like to thank Dr. Maria Genaleen Q. Diaz for her technical expertise and valuable insights throughout this research. We would also like to express our sincere appreciation to the Plant Cell and Tissue Culture Laboratory (PCTC) Laboratory and National Plant Genetic Resources Laboratory (NPGRL) of the Institute of Plant Breeding (IPB), College of Agriculture and Food Science (CAFS), UPLB for providing us with plant materials for banana accession ‘Mapilak’. Additionally, sincere appreciation is directed towards Mr. Rodelio R. Pia, Mr. Ronilo R. Bajaro, and Garee R. Hernandez for their invaluable contributions in providing technical assistance.

## AUTHOR APPROVALS

All authors reviewed, revised, and approved the final version of the manuscript.

## DATA AVAILABILITY

The dataset(s) generated in this study are included within the article and Supplemental file(s) are available from the corresponding author on practical request.

## ETHICS APPROVAL

Not applicable.

## CONSENT TO PARTICIPATE

Not applicable.

## CONSENT TO PUBLISH

Not applicable.

